# The hemodynamic signal as a first-order low-pass temporal filter: Evidence and implications for neuroimaging studies

**DOI:** 10.1101/077859

**Authors:** Antoine Sauvage, Guillaume Hubert, Jonathan Touboul, Jérôme Ribot

**Author notes:** Equal contribution (Jonathan Touboul, Jérôme Ribot).

## Abstract

Neuronal activation triggers blood flow and hemoglobin deoxygenation. These hemodynamic signals can be recorded through magnetic resonance or optical imaging, and allows inferring neural activity in response to stimuli. These techniques are widely used to uncover functional brain architectures. However, their accuracy suffers from distortions inherent to hemodynamic responses and noise. The analysis of these signals currently relies on models of impulse hemodynamic responses to brief stimuli. Here, in order to infer precise functional architectures, we focused on integrated signals associated to the dynamical response of functional maps. To this end, we recorded orientation and direction maps in cat primary visual cortex and confronted two protocols: the conventional episodic stimulation technique and a continuous, periodic stimulation paradigm. Conventional methods show that the dynamics of activation and deactivation of the functional maps follows a linear first-order differential equation representing a low-pass filter. Comparison with the periodic stimulation methods confirmed this observation: the phase shifts and magnitude attenuations extracted at various frequencies were consistent with a low-pass filter with a 5 s time constant. This dynamics open new avenues in the analysis of neuroimaging data that differs from common methods based on the hemodynamic response function. In particular, we demonstrate that inverting this first-order low-pass filter minimized the distortions of the signal and enabled a much faster and accurate reconstruction of functional maps.

## 1 INTRODUCTION

Neuroimaging techniques have rapidly developed in the last decades, allowing visualizing *in vivo* neural activity over large cortical regions with fine spatial and temporal resolutions. One major difficulty however lies in the multiple sources of noise that distort the recorded signal. This greatly limits the efficiency of the technique and often requires multiple repetitions of stimuli, increasing substantially the experiments duration and thus limiting the number of conditions tested. Improving signal to noise ratios and rapidity of acquisition has thus been a great endeavor in neuroimaging and the topic of important research.

Among the different neuroimaging techniques, those based on the hemodynamic signals, such as functional magnetic resonance imaging (fMRI) or intrinsic optical imaging, have led to major discoveries in neuroscience. The study of these dynamics has opened the way to the development of analysis techniques for deciphering the underlying neuronal responses. Ample theoretical and experimental work has been devoted to the fine characterization of the hemodynamic response function (HRF) following a short stimulus presentation (Glover, 1999) and their relationship with increased metabolism associated to neurons activation. Indeed, neural activation triggers blood flow and venous vessels dilation to meet the needs in oxygen and glucose to support neuron function (Buxton, 1998; Vanzetta *et al.*, 2004; Vanzetta and Grinvald, 2008). The question of how basic hemodynamic responses are combined in response to complex stimuli has attracted much attention. The resulting models are generally mathematically involved, but efficiently capture the main characteristics of the response dynamics (see e.g. Friston et al., 2000). These models have been particularly used in fMRI studies and have led to important advance in the understanding of brain function (see Lindquist et al., 2008 for an overview of applications).

HRF has been recently used to analyze functional imaging signals resulting from the periodic repetition of brief stimuli (See e.g. Arnett et al., 2014; Sun & al., 2013; Chen-Bee et al, 2010). By carefully choosing the time interval between each repetition, the resulting hemodynamic signal behaves like a sine function that can be readily analyzed through Fourier transform. The hemodynamic delay is then subtracted from the signal, and the correspondence with the stimulus is achieved. This paradigm has allowed to drastically reduce the recording time while improving the signal to noise ratio. However, this technique still suffers from the relatively limited number of conditions that can be tested during a single experiment. In order to increase the number of stimulus conditions, a protocol based on a continuous activation of the neurons was proposed (See e.g. Yakoub et al, 2008; Vanni et al. 2010a; Ribot et al. 2013, 2016). This protocol relies on the uninterrupted and periodic presentation of a full set of parameters

Although the latter stimulation technique is theoretically more powerful, the recorded signal faces additional challenges. First, the complex, non-linear transformation of the underlying neural activity through the hemodynamic response (See Soltysik et al., 2004 for a review) makes the signal interpretation strenuous. Moreover, adaptation mechanisms and phase inconsistency over such long, continuous periods of stimulation may impact recordings of neural activity (Sun & al., 2013). These effects may thus considerably degrade the signal interpretation, and may be the cause of important inconsistencies. For instance, a recent experiment on cat primary visual cortex using this technique underlined that thus obtained direction map only mildly exhibits the fundamental properties observed in those resulting from conventional techniques (Vanni et al., 2010a).

The present paper undertakes a fine investigation of the filtering properties of hemodynamic signals when recorded with the continuous, periodic stimulation protocol and confronts these responses to conventional, episodic techniques. By considering specific collective statistics of the hemodynamic response, we show that filtering properties average out into a simple model that reduces to a first order low-pass filter. To this purpose, we collected and analyzed intrinsic optical imaging data in cat early visual cortex. The first step to understand these effects was to isolate the dynamics of the hemodynamic signal. To this purpose, we show that, in response to stimuli with various durations, the magnitude of the functional maps evolves as the response of a first-order linear dynamical system to the stimulus presentation, and thus acts as a low-pass temporal filter. This observation led to predict specific phase shifts and signal attenuations in response to continuous, periodic stimulations; these depend on (i) the harmonic of the signal considered and (ii) on the stimulation frequency. We tested that prediction and found that the resulting Bode diagrams coincided with the estimated low-pass filter. These distortions being finely estimated, we recovered accurate and reliable functional maps by simply inverting the estimated filter. We used this technique to obtain very fine direction maps in cat, and recover a precise orthogonal relationship with the orientation map.

## 2 MATERIALS AND METHODS

Five adult cats (two male, three females), born from different litters in our colony, were studied. They were all in good health and had no apparent malformations or pathologies. All experiments were performed in accordance with the relevant institutional and national guidelines and regulations i.e. those of the Collège de France, the CNRS and the DDPP (JO 87-848, consolidated after revision on May 30, 2001, Certificates n°^s^ 75-337, 75-1753, 75-1754, French “Ministère de l’Agriculture et de la Pêche”). They also conformed to the relevant regulatory standards recommended by the European Community Directive (Directive 2010/63/UE) and the US National Institutes of Health Guidelines.

### 2.1 Surgical procedure

On the day of the optical imaging experiment, animals were anaesthetized with Saffan® (initial, i.m., 1.2 mg/Kg; supplements, 1:1 in saline, i.v. as needed). After tracheal and venous cannulation, electrocardiogram, temperature and expired CO_2_ probes were placed for continuous monitoring. Animals were installed in the Horsley-Clarke stereotaxic frame and prepared for acute recordings. The scalp was incised in the sagittal plane, and a large craniotomy was performed overlying areas 17 and 18 of both hemispheres. The nictitating membranes were then retracted with eye drops (Neosynephrine® 5%, Ciba Vision Ophthalmics, France) and the pupils dilated with atropine eye drops (Atropine 1%, MSD-Chibret, France). Scleral lenses were placed to protect the cornea and focus the eyes on the tangent screen 28.5 cm distant. The size of the lenses was adapted to the eye of each cat. Animals were then paralyzed with an infusion of Pavulon (0.2 ml/kg i.e. 0.4 mg/kg i.v.) and breathing was assisted artificially through a tracheal cannula with a 3:2 mixture of N_2_O and O_2_ containing 0.5–1.0% isoflurane. The respiration frequency was adjusted to around 18 per minute and the volume adapted to the ratio of exhaled CO_2_ (pCO_2_ was maintained at 4%). Paralysis was maintained throughout the recording by continuous infusion of a mixture of Pavulon (0.1 ml/kg/h) diluted in glucose (5%) and NaCl (0.9 g/l). At the end of a recording session, the animal was given a lethal dose of pentobarbital. The experiments lasted in total less than 12 hours.

### 2.2 Optical imaging

The cortex was illuminated at 545 nm to reveal the vascular pattern of the cortical surface and at 700 nm to record the intrinsic signals. The focal plane was adjusted to 500 µm below the cortical surface. The optic discs were plotted by tapetal reflection on a sheet of paper covering the tangent screen placed 28.5 cm in front of the animal. The center of the screen was situated 8 cm (∼15°) below the middle of the two optic discs, located at approximately at 8.5° into the lower visual field of the animal (Bishop et al., 1962). Intrinsic optical signals were recorded while the animals were exposed to visual stimuli displayed on a CRT monitor. The size of the display subtended a visual angle of ∼75° × 56°. The image had a resolution of 14 pixels per degree and the refresh rate was 88 Hz.

Two stimulation protocols were used: the episodic one and the continuous one (see details below). Frames were acquired by CCD video camera (1M60, Dalsa, Colorado Springs, USA) at the rate of 40 frames per second. In the episodic protocol, frames were then stored after binning by 2 × 2 pixels spatially and by 40 or 8 frames temporally using the VDaq system (Optical Imaging Inc., New York, USA). In the continuous stimulation protocol, frames were stored after binning by 2 × 2 pixels spatially and by 12 frames temporally using the LongDaq system (Optical Imaging Inc., New York, USA). Images were acquired with a resolution of approximately 25 µm/pixel, or by using a teleconverter with a two-fold finer resolution.

### 2.3 Episodic stimulation protocol

#### Stimulation

In the episodic stimulation protocol, the animal was stimulated with full screen sine-wave gratings with a spatial frequency of 0.27 cycle/degree. The gratings were drifting at 24 equally spaced directions of motion. The temporal frequency of the gratings was fixed at 2 Hz. Each stimulus was presented once in a pseudo-random sequence in a single trial of recordings, and interleaved with blank-screen presentation for 15 s. Fifty trials were collected in each recording session. The intrinsic optical signal in response to each stimulus was recorded immediately after the stimulus onset. A total of five frames of 1s duration each were recorded. Only the last 3 frames, which correspond to the strongest intrinsic signal, were used for data analysis.

#### Data analysis

Data were analyzed with a multivariate analysis technique, the generalized indicator function method (Yokoo et al., 2001; Ribot et al., 2006). This method identifies the stimulus-related activity patterns (signals), without prior knowledge about the signal and noise characteristics. The extracted signals, that is, the generalized indicator functions, are determined by maximizing the weighted difference between the signal variance and the noise variance. This algorithm efficiently extracts stimulus-related activity patterns from noisy signals originating mainly from blood vessels and spatially slowly varying fluctuations inherent to the recorded signals. Finally, a Fourier transform was performed on the signals for each pixel and the phase and magnitude at several harmonics were extracted.

#### Calculation of the direction map

To calculate the direction map (section 3.3), the data obtained with the periodic stimulation protocol was first shifted so that the minimum value of the response at each pixel is equal zero, and then interpolated by the sum of two Von Mises function (Swindale et al., 2003):

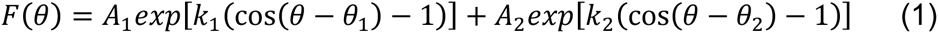

where *θ* is the direction of stimulus motion, *θ*_*1*_ and *θ*_*2*_ represent the preferred and null directions, *k*_*1*_, *k*_*2*_ are inversely related to the tuning width of each peak and *A*_*1*_, *A*_*2*_ are their corresponding amplitudes (*A*_*1*_ > *A*_*2*_). For each response obtained at each pixel, we used the Levenburg-Marquardt optimization algorithm to determine the parameters *θ*_*1*_, *θ*_*2*_, *k*_*1*_, *k*_*2*_, *A*_*1*_ and *A*_*2*_ that minimize the residual sum of squares. This optimization algorithm is iterative and requires the definition of an initial parameter set, which was simply chosen from the response curve as the positions (*θ*_*1*_, *θ*_*2*_) and amplitudes (A_1_, A_2_) of the two maximas, while *k*_*1*_ and *k*_*2*_ were set to 1. The programs were written in Python and used the spicy.optimization module (Jones et al, 2001).

#### Dynamics of the orientation magnitude

For the dynamics of orientation magnitudes of Fig. 1, the response to gratings drifting at eight different directions of motion was recorded for 20 s each at a frame rate of 5 Hz. Stimuli were presented for 1s, 5s or 10s. Data were processed frame by frame with the indicator function method and vector summed to produce orientation maps.

**Figure 1.**
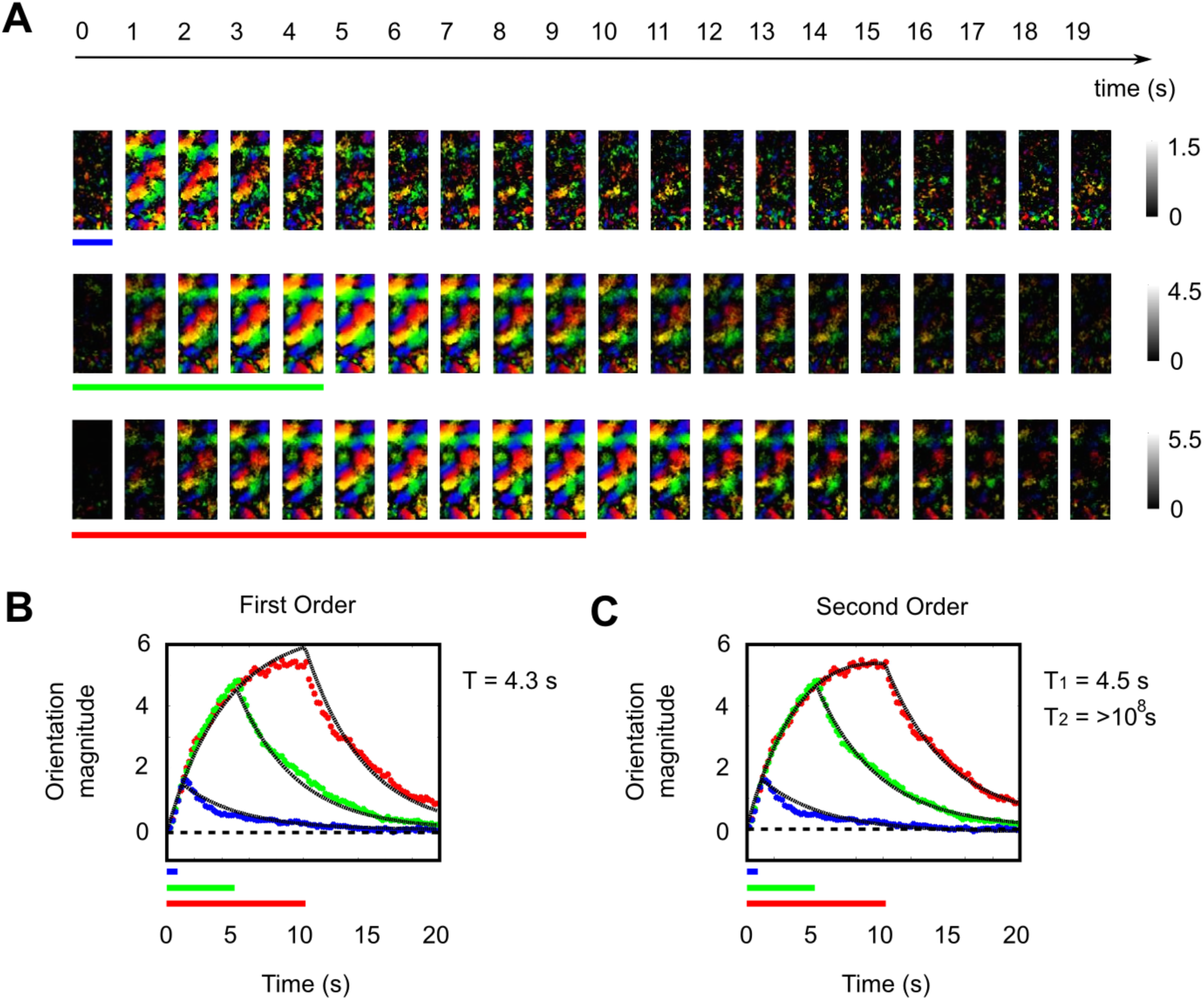
Orientation magnitude evolves as a first-order temporal low-pass filter. (A) Orientation polar maps recorded with conventional techniques when the animal is stimulated with gratings for durations of 1 s (top), 5 s (middle), and 10 s (bottom). The total recording time after stimulus onset is 20s. Signals were recorded every 0.2 s but only data obtained every 1 s are shown. Note that the scale bar was adapted for each condition. (B) The total orientation magnitude of the maps shown in (A) was fitted with a first-order low-pass filter (black curve) for the three stimulation durations. (C) Same as (B) for a second-order low-pass filter.

### 2.4 Continuous stimulation protocol

#### Stimulation

In the continuous stimulation protocol, full-screen visual stimuli were presented continuously to the animal. Each stimulus consisted of sine-wave gratings drifting in one direction and rotating in a counter-clockwise manner (Kalatsky and Stryker, 2003). The temporal frequency of the drifting was set at 2 Hz. In some animals, only one spatial frequency was presented, usually 0.27 cycle/degree. In other cases, 12 or 30 spatial frequencies ranging linearly in a logarithmic scale were presented in random order. The angular speed of the rotation was 2 rotations per min. In one experiment, several angular speeds were used ranging between 6s and 120 s. For each stimulus condition, the duration of the recording was around 5 min.

#### Image processing

Data analysis was based on the method developed by Kalatsky and Stryker (2003) to extract periodic signals from intrinsic data in Fourier space. First, the generalized indicator function method (Yokoo et al., 2001; Ribot et al., 2006) was applied to raw data for each stimulus condition separately. Then, a Fourier transform was performed on the temporal signal of each pixel and the phase and magnitude at several harmonics were extracted.

#### Calculation of the direction map

In order to calculate the direction maps, we first pre-processed our data by inverting the first-order low-pass filter with the adequate τ value (see section 3.2.3.). Phase and magnitude maps were then extracted at *H*_*1*_ and *H*_*2*_ using the Fourier transform. Then, we reconstructed at each pixel the response tuning curve by summing up signals extracted at *H*_*1*_ and *H*_*2*_. The preferred direction was defined as the stimulus direction that reached the maximal response.

### 2.5 Analysis of phase maps

#### Autocorrelation

The periodic structure of the phase map was investigated by calculating the 1D-autocorrelation function over 2 mm of the cortex. The 2D autocorrelation map was first calculated, and values were averaged radially to obtain the 1D profile.

#### Comparison between phase maps

To compare phase maps obtained in the two protocols, phase maps for each harmonic were subtracted from each other and circularly averaged. The resulting angle was defined as the average phase shift, and the resulting norm was defined as the correlation index (between 0 and 1).

## 3 RESULTS

### 3.1 THE HEMODYNAMIC SIGNAL AS A FIRST-ORDER TEMPORAL FILTER: EVIDENCE AND IMPLICATIONS

The first step in our study consisted in characterizing the dynamical response of the hemodynamic signal in the episodic stimulation protocol, and its consequences on the signals obtained with continuous periodic stimulation protocols.

#### 3.1.1 Dynamics of the functional maps with the episodic stimulation

In the first experiment, we used the conventional, episodic stimulation method (see Methods) to reconstruct the orientation map in cat visual cortex with intrinsic optical imaging. Oriented, drifting gratings were presented to the animal for various durations *T*_*stim*_ = 1 s, 5 s and 10 s. Intrinsic signals were recorded for 20 s after the stimulus onset. Orientation polar maps for each stimulus duration are shown in Fig. 1A, and the time evolution of the total orientation magnitude in Fig. 1B-C. In accordance with previous results (Grinvald et al., 1999), we observed that the magnitude increased shortly after stimulus presentation, peaked around the end of the stimulation time, and then decreased slowly to the baseline level.

These time courses reflect the low-pass properties of the hemodynamic signal and provide an estimate of the actual parameters that describe its dynamics. Mathematically, these dynamics are evocative of first order linear differential equations:

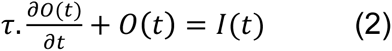

where *t* represents time, τ is the time-constant of the system, *O(t)* is the output signal triggered by the input presentation *I(t)*. The general solution of the equation is given by:

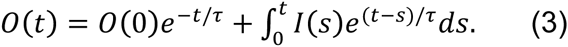

We represent in Fig. 1B the good agreement obtained by fitting this dynamics to the data obtained by presenting a constant stimulus for a duration *T*_*stim*_:

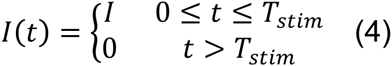

The solution is thus given by

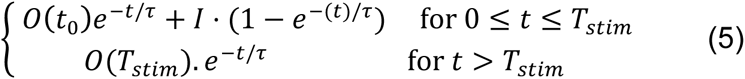

The best fit of the time course for all stimulus durations together was for τ = 4.3 *s*. This fit was close to optimal for a 5 s stimulus duration (green curve, linear regression, R^2^ = 0.98) and for a 10 s stimulus duration (blue curve, R^2^ = 0.97), but slightly deviated the actual time courses for a 1 s stimulus duration (red curve, R^2^ = 0.92).

First order differential equation systems are very simple models that may ignore important phenomena arising at different timescales. We thus tested more complex models in order to reveal other important phenomena beyond the first-order low-pass filtering properties. We found that higher order differential equations that fitted best the data had degenerate higher order differential terms, and were effectively first order models in disguise. For instance, second order differential equation (Fig. 1C) modeling signals as combinations of two exponentials with distinct timescales τ_1_ and τ_2_ had one degenerate exponential (τ_1_ = 4.5 *s* and τ_2_ > 10^8^ *s*), showing that the dynamics is effectively of order 1.

#### 3.1.2 Implications for neuroimaging studies

The amplitude-phase Bode plot for this first-order low-pass filter is shown in Fig. 2A-B. The common stimulation frequency range used in previous studies is shown in gray, and the arrow locates the cutoff frequency ω_*c*_ = (*2*πτ)^−1^ (≈ 1/27 Hz), which falls within the stimulation range. These diagrams indicate that for harmonics extracted around the period of stimulation, the magnitude between the input and the output signals should be slightly decreased, and the phase difference should be shifted by several dozen degrees. For harmonics extracted at higher frequencies, the attenuation and phase shift should be even more dramatic.

**Figure 2.**
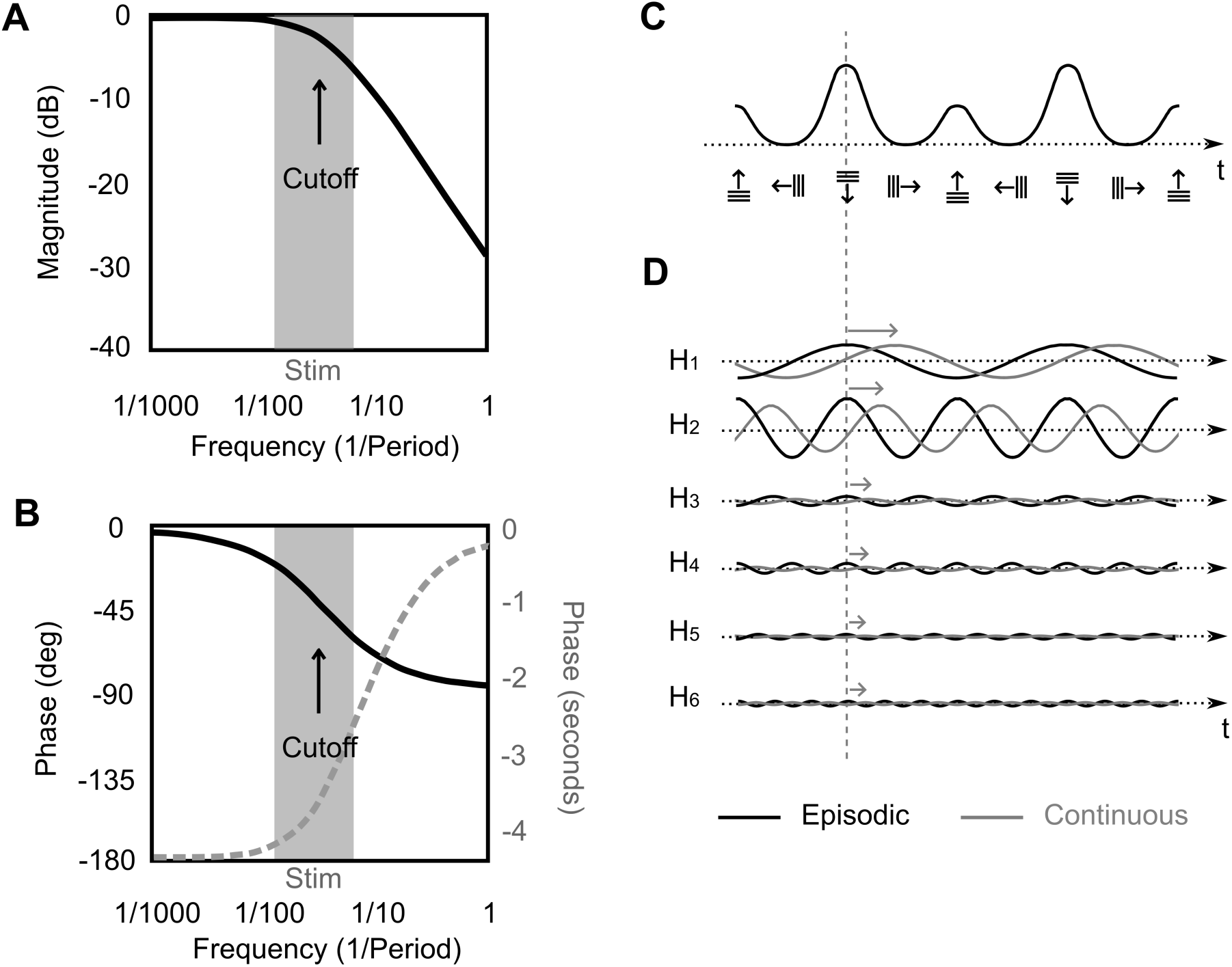
Impact of the low-pass nature of the hemodynamic signal on extracted harmonics. (A-B) Bode plot of the magnitude (A) and phase (B) for the first-order low-pass filter found in Fig. 1. The gray area corresponds to the frequency range typically used for the stimuli. Note that higher harmonics can fall on the right side of this zone. The cutoff frequency found in Fig. 1 is indicated with an arrow. We indicated in (B) the phase Bode plot expressed in seconds in gray dashed line. (C) Schematic tuning curve of a population of neurons in one cortical column when recorded with classical, conventional techniques. In this example, the preferred orientation of the neurons is horizontal and their preferred direction is downwards. (D) Schematic representation of signals extracted in the case of the episodic stimulation method (black curves) and of the continuous stimulation protocol (gray curves). The first six harmonics are depicted. The impact of the phase shift shown in (B) is depicted with gray arrows.

As shown in Fig. 2C-D, this should greatly impact the hemodynamic signal when rotating gratings are used to extract functional maps in the primary visual cortex. Figure 2C sketches a typical activity that should be recorded with classical, episodic techniques at a location in the visual cortex. The signal peaks at the preferred direction of the recorded neurons, has a secondary peak around 180° from the preferred direction, and is minimal at orthogonal orientations. Signals extracted at the six first harmonics are depicted in Fig. 2D, black curves. For continuous stimulation paradigms, the stimulus consists in oriented gratings, drifting in one direction, and rotating at fixed angular value. Fourier transform is applied at each pixel (Fig. 2D, gray curves), and the information contained at various harmonics of the frequency of rotation is extracted. Since different phase shifts however occurs depending on the extracted frequency (Fig. 2A), different phase delays (gray arrows) should be observed in the signal with increasing harmonics. Moreover, these phase shifts should also vary depending on the actual stimulation cycle value, and we provide further evidence in the following section.

### 3.2 THE PERIODIC STIMULATION APPLIED TO CAT VISUAL CORTEX

#### 3.2.1 Visualization of the phase maps

We recorded on the same animal the hemodynamic signals with the continuous, periodic stimulation protocol (rotation cycle equal to 30 s). The cumulative power spectrum over the region of interest is shown in Fig. 3A. Three main peaks emerged, corresponding to the frequency of rotation *H*_*1*_ and the its two harmonics *H*_*2*_ and *H*_*4*_.

**Figure 3.**
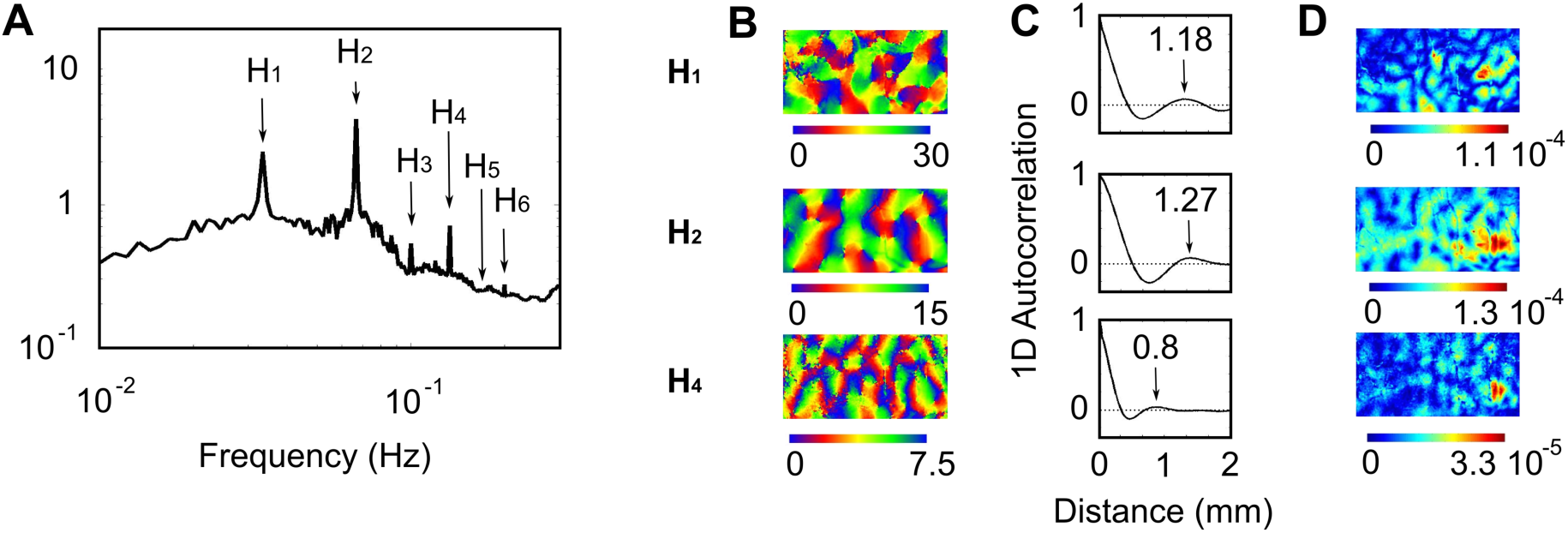
Visualization of the phase maps extracted at three dominant harmonics. (A) Cumulative power spectrum over a region of interest covering both area 17 and area 18. The continuous stimulation protocol was used. The peak corresponding to the frequency of rotation (*H*_*1*_ = 1/30s) as well as those corresponding to other harmonics (*H*_*2*_ through *H*_*6*_) are indicated with an arrow. (B) Phase maps for the harmonics *H*_*1*_, *H*_*2*_ and *H*_*4*_. Scale bar is in seconds. The corresponding 1D-autocorrelograms are shown in (C). The periodicity is indicated with an arrow. The magnitude map for each phase map is shown in (D).

The phase maps for these three frequencies are shown in Fig. 3B: that of *H*_*2*_ is reportedly related to the preferred orientation map (Kalatsky and Stryker, 2003; Zepeda et al., 2004; Jha et al., 2005; Yakoub et al., 2008; Vanni et al., 2010a), and that of *H*_*1*_ to the preferred direction map (Vanni et al., 2010a). They both contain a periodic structure, as indicated by the peak in the 1D autocorrelation in Fig. 3C (black arrow), and have comparable magnitude (Fig. 3D).

The magnitude of the phase map of the fourth harmonic *H*_*4*_ (Fig. 3D) is much weaker than those found for *H*_*1*_ and *H*_*2*_, arguing for a minor role in the total signal. However, this phase map (Fig. 3B) contains a periodic structure (Fig. 3C). Moreover, this signal is biologically interpretable: it reflects changes in orientation tuning width. We will thus also consider this harmonic in the rest of our analyses.

#### 3.2.2 Phase shifts are observed during continuous stimulation protocols

We compared the three phase maps obtained with conventional techniques based on episodic stimulation (Fig. 4A, see single condition maps on Suppl. Info. 1) with those obtained with the periodic stimulation protocol (Fig. 4B, top panels, see single condition maps on Suppl. Info. 1). The angular rotation frequency of the gratings was 1/30 Hz, a value close to the actual cut-off frequency found in Fig. 1 (ω_*c*_ ≈ 1/27 Hz). The distribution of phase differences between the two paradigms are shown in Fig. 4B, bottom panels (phase maps obtained with the episodic stimulation protocol were rescaled in seconds to allow for comparison). For each set of phase maps, a clear peak could be observed (black arrows), indicating a correlation between them. However, both the height and the position of the peak varied for the different harmonics.

**Figure 4.**
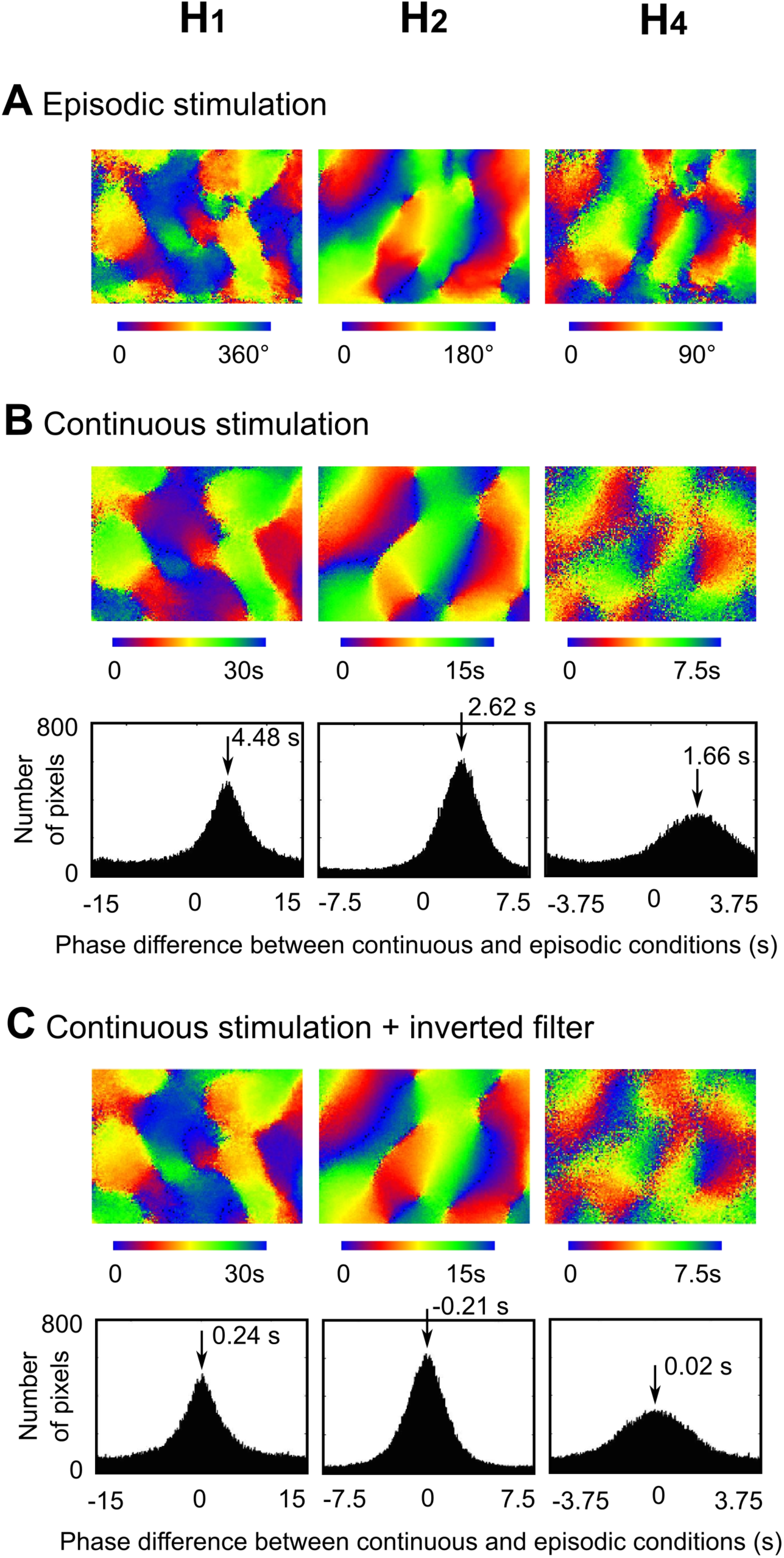
The phase delay varies for each extracted harmonics. (A) Phase maps obtained with the episodic presentation of the stimuli. Scale bar is in degree. (B) Top: Phase maps for *H*_*1*_, *H*_*2*_ and *H*_*4*_ obtained with the continuous presentation of orientated, drifting gratings rotating at the frequency of 1/30s. Scale bar is in seconds. Bottom: Histograms showing the difference between phase maps obtained with the two protocols. For simplicity, the scale bar in (A) was adapted to the one in (B). (C) Same as (B) when data where pre-processed by inverting the first-order low-pass filter (τ = 5.83 *s*). Single condition maps for phase maps in (A) and (B) are shown in Supplementary Info. 1.

These differences were quantitatively investigated in Table 1 (N = 4 animals). We found that maps related to *H_2_* had the best correlation index, particularly for domains with the highest magnitude (t-test, p<0.0001). The delay between the two methods for this harmonic was around 2.4 s and was independent of the magnitude of the signal (t-test, p>0.84). The correlation between maps for the two other harmonics was weaker than this for *H_2_* (t-test, p<0.0001 and p=0.0002 for *H*_*1*_ and *H*_*4*_ respectively). But this correlation increased significantly when solely domains with the highest magnitude were kept (t-test, p<0.0001 and p=0.0009 for *H*_*1*_ and *H*_*4*_ respectively). The phase delay between the two methods was maximal for *H*_*1*_ (around 4.4 s, t-test, p<0.0001 against H_2_ and H_4_) and minimal for *H*_*4*_ (around 1.5 s, t-test, p<0.0001 against H_1_ and p=0.0032 against H_2_). However, no difference in phase delay could be found when only domains with the highest magnitude were considered (t-test p>0.87 and p>0.54 for H_2_ and H_4_ respectively).

**Table 1.**
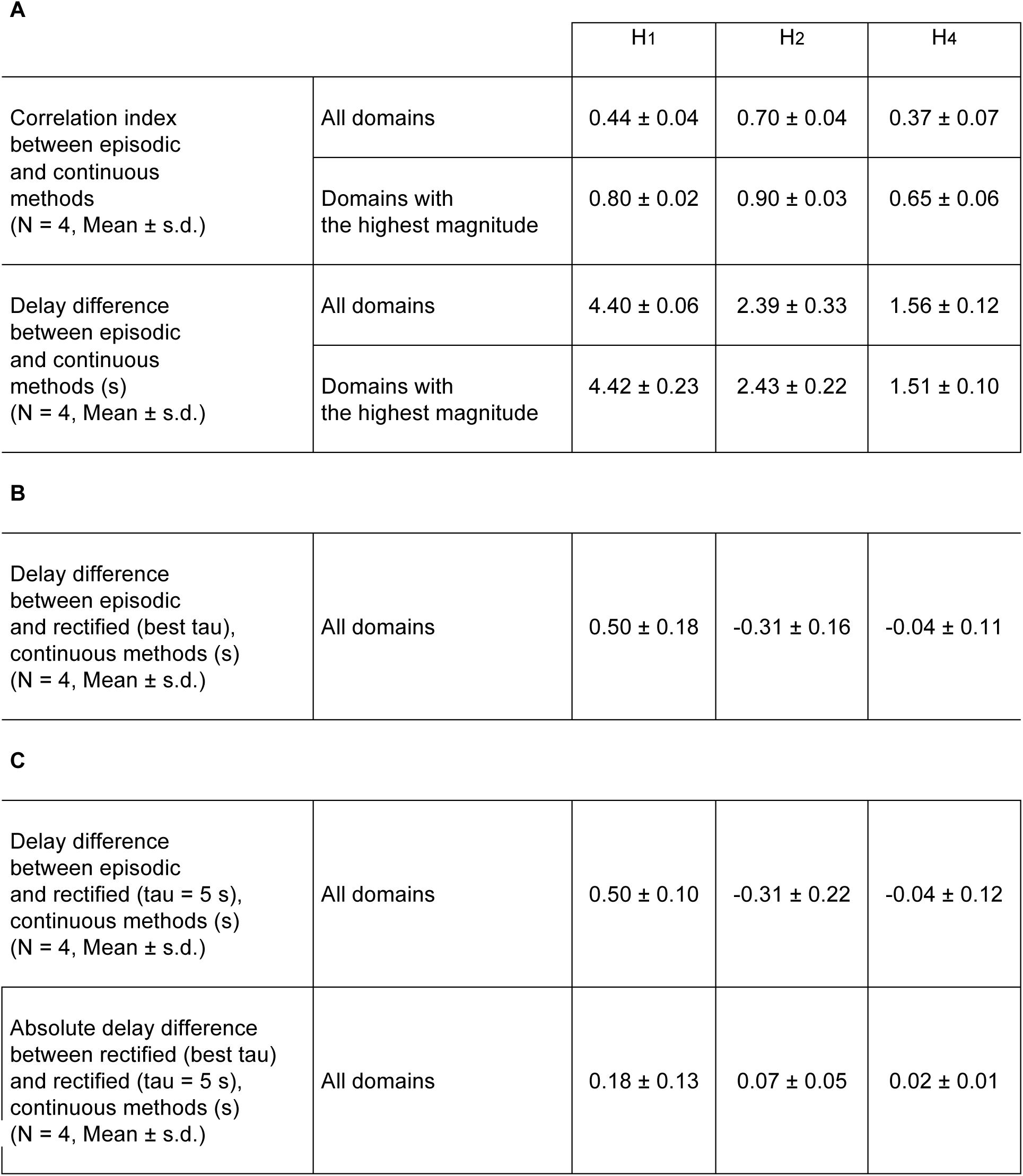
Phase shifts and correlation indices between episodic and periodic paradigms. (A) For the three main harmonics *H*_*1*_, *H*_*2*_ and *H*_*4*_, the average values (± s.d.) over the region of interest of the correlation index and of the delay shift are shown. These values were also calculated over cortical domains with the highest magnitude. (B) Average (± s.d.) delay shift calculated after inverting the first-order low-pass filter with the best τ value for each animal. (C) Average (± s.d.) delay shift calculated after inverting the first-order low-pass filter with τ = 5 *s*. For each animal, we also calculated the absolute error made when taking the best τ value and τ = 5 *s*. Average error (± s.d.) is shown.

|  |  | H1 | H2 | H4 |
| --- | --- | --- | --- | --- |
| Correlation index between episodic and continuous methods<br>(N = 4, Mean ± s.d.) | All domains | 0.44 ± 0.04 | 0.70 ± 0.04 | 0.37 ± 0.07 |
| | Domains with the highest magnitude | 0.80 ± 0.02 | 0.90 ± 0.03 | 0.65 ± 0.06 |
| Delay difference between episodic and continuous methods (s)<br>(N = 4, Mean ± s.d.) | All domains | 4.40 ± 0.06 | 2.39 ± 0.33 | 1.56 ± 0.12 |
| | Domains with the highest magnitude | 4.42 ± 0.23 | 2.43 ± 0.22 | 1.51 ± 0.10 |
| **B** |  |  |  |  |
| Delay difference between episodic and rectified (best tau), continuous methods (s)<br>(N = 4, Mean ± s.d.) | All domains | 0.50 ± 0.18 | -0.31 ± 0.16 | -0.04 ± 0.11 |
| **C** |  |  |  |  |
| Delay difference between episodic and rectified (tau = 5 s), continuous methods (s)<br>(N = 4, Mean ± s.d.) | All domains | 0.50 ± 0.10 | -0.31 ± 0.22 | -0.04 ± 0.12 |
| Absolute delay difference between rectified (best tau) and rectified (tau = 5 s), continuous methods (s)<br>(N = 4, Mean ± s.d.) | All domains | 0.18 ± 0.13 | 0.07 ± 0.05 | 0.02 ± 0.01 |

These results therefore confirm our hypothesis that the low-pass filter’s properties of the hemodynamic signal induce changes in the phase of the extracted harmonics. Moreover, since they do not differ between domains of the highest and the lowest magnitude, they are independent on the actual neural properties of cortical locations.

#### 3.2.3 Phase maps are recovered by inverting the temporal low-pass filter

The observed phase differences are quantitatively important, since they correspond to shifts of around 53°, 29° and 18° in the phase maps of *H*_*1*_, *H*_*2*_ and *H*_*4*_ obtained with conventional techniques (Fig. 4A). A rectification is thus necessary to recover the actual functional maps as obtained with conventional episodic methods. We show in this section that deconvolving the recorded signal with a first-order temporal filter provides a simple and efficient solution to this issue.

It is particularly convenient to invert the filter working in Fourier space. Indeed, in this space, the hemodynamic filter is written as

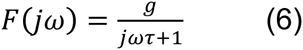

where *j* ^2^ = −1, ω is the frequency, τ is the filter time constant and *g* is the gain of the filter. The phase shift induced by this filter is:

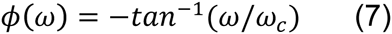

where ω_*c*_ = *2*πτ ^−1^ is the cutoff frequency.

In Fig. 4, the best fit of the phase shift with biological data was for τ = 5.83 *s* (R^2^ = 0.96). Pre-processing the data with the inverted filter

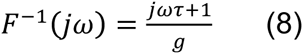

should thus improve the reconstruction of the phase maps. We tested this technique, fixing the gain *g* equal to 1. We compared the rectified phase maps (Fig. 4C) to those obtained the episodic stimulation and confirmed the drastic reduction of the phase shift compared to the non-rectified map, with absolute values less than 0.25 s. The shifts corresponded to deviations around 2.9°, −2.6° and 0.2° between the phases maps obtained with continuous and episodic techniques. This shift was found highly significant for the four animals tested (Table 1B, t-test, p<0.0001 for each harmonic), although a residual delay shift remained, especially for *H*_*1*_ and *H*_*2*_.

Remarkably, we found a high consistency in the time constant τ defined for each animal: Values ranged between 4.66 s and 5.83 s, with a mean value of 5.03 ± 0.55 s.d.). Instead of adapting τ precisely for each animal, this suggests that this value can readily be used for inverting the low-pass filter. We tested this hypothesis in Table 1C by taking τ = 5 *s*. This value was sufficient for recovering the residual phase shifts shown in Table 1B. The average absolute error remained small for each animal: 0.18 s for *H*_*1*_ (2.2 deg), 0.07 s for *H*_*2*_ (0.8 deg) and 0.02 s for *H*_*4*_ (0.2 deg). We conclude that the actual phase maps obtained with time-consuming, episodic techniques can be efficiently recovered from fast, continuous, periodic stimulation protocols by inverting the low-pass filter with τ = 5 *s*.

#### 3.2.4 Variations of the phase delays with rotation cycle

As indicated section 3.1.2, the phase shifts should also vary with the stimulus frequency. In order to test this dependence, we report in Fig. 5 the result of an experiment consisting in stimulating the animal with gratings rotating at various speeds (period of rotation ranging from 6 s to 120 s). Figure 5A shows the maps extracted at *H*_*2*_ for each condition. These maps were compared to the one obtained during an episodic presentation of the gratings (Fig. 5B). Figure 5C (green dots) shows the average phase difference for this harmonic between the two protocols. It was minimal for a period of rotation of 6 s (around 0.9 s delay), increased logarithmically with increasing period of rotation (Linear regression, R^2^ = 0.65), and was maximal for 120 s (around 5.3 s delay). This linear increase was also present in the two other harmonics *H*_*1*_, and *H*_*4*_ (blue and red dots, respectively, R^2^ = 0.69 and 0.67), but corresponded to different slopes; as in Fig. 4, the phase shift for *H*_*1*_ was larger than that of *H*_*2*_ while the phase shift of *H*_*4*_ smaller.

**Figure 5.**
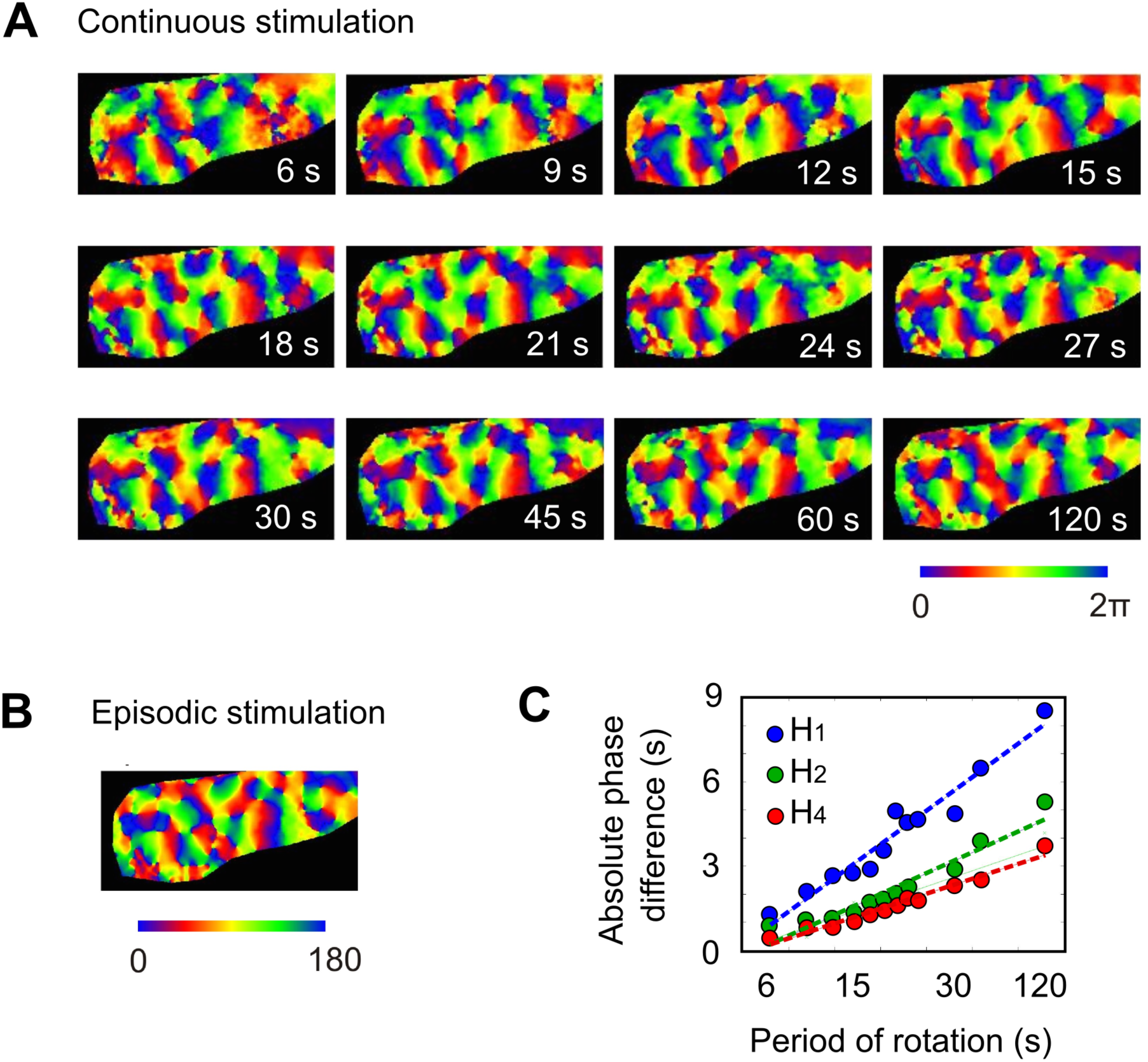
The phase shift varies with the frequency of rotation. (A) Representation of the phase maps extracted at *H*_*2*_ for various periods of rotation (6, 9, 12, 15, 18, 21, 24, 27, 30, 45, 60 and 120 seconds). (B) Orientation map obtained with the episodic presentation of the stimuli. (C) Phase shift between the two conditions are shown with respect to the periods of rotation. X-scale is logarithmic.

#### 3.2.5 The magnitude Bode plot

The Bode plot for magnitude for the same data as in Fig. 5 is represented in Fig. 6A, superimposed in black, full line, to the first-order magnitude Bode plot (τ = 5 *s*, *g* = 1). The agreement with biological data was acceptable (Linear regression, R^2^ = 0.735), but could be increased by adapting the value of the gain. As shown in Fig. 6B, the mean square error was minimal for a gain equal to 0.92. For this value, the coefficient of determination R^2^ increased to 0.796 (Fig. 6A, black dashed line). For the highest frequencies > ω_*c*_, the fit with biological data remained high (Linear regression, R^2^ = 0.734), indicating a good agreement with the 20 dB/decade magnitude decrease typical of a first-order low-pass filter.

**Figure 6.**
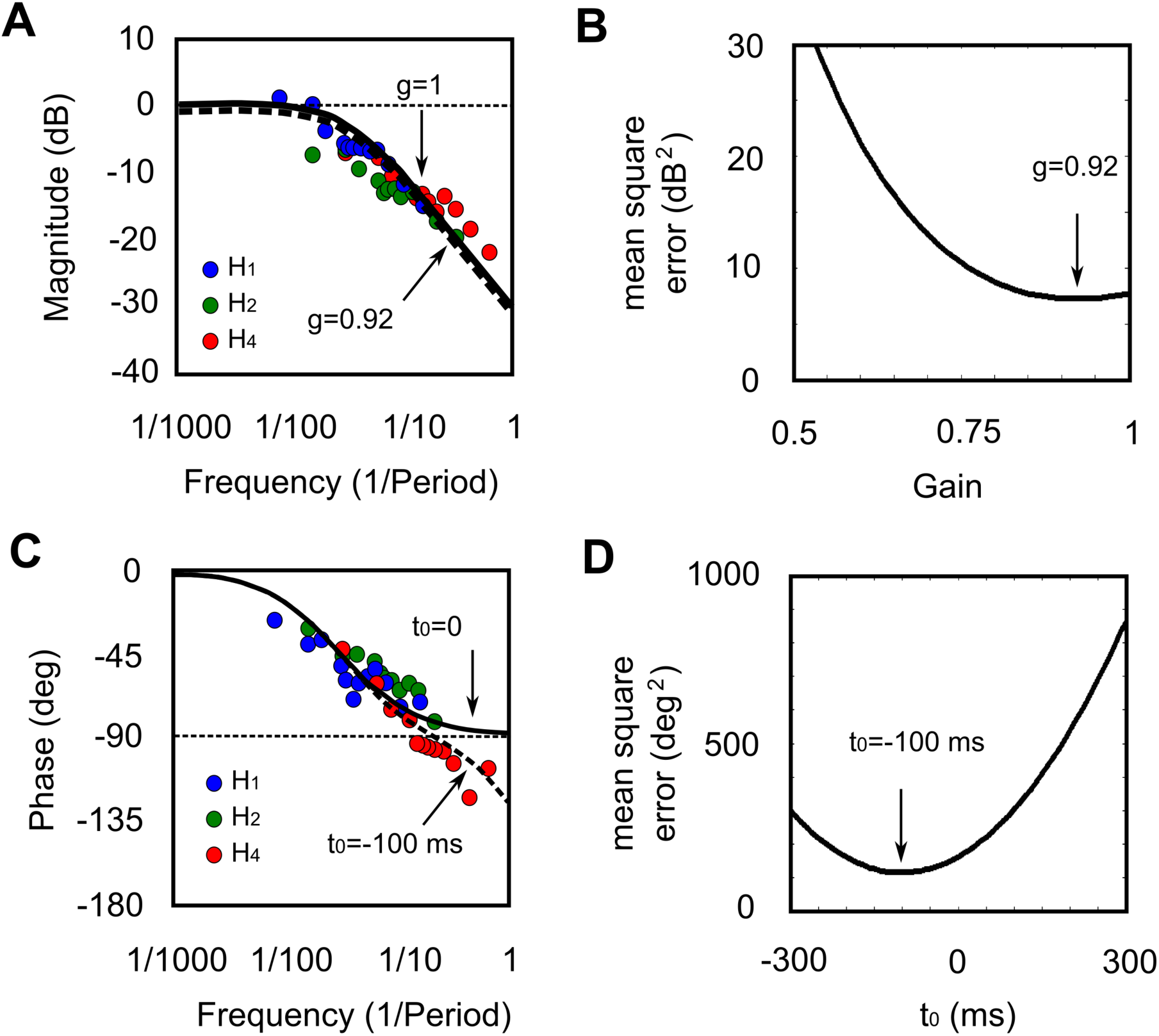
Tuning gain and delay. (A) Bode plot of the magnitude. The black full curve corresponds to the Bode plot with τ = 5 *s* for a gain equal to 1, and the dashed curve is the one for a gain equal to 0.82. (B) Mean square error between the data and the model for different values of the gain g. (C) Bode plot of the phase for each rotation period and each harmonic. The solid curve corresponds to the Bode plot with τ = 5 *s*, and the dashed curve is obtained when considering a constant delay *t*_0_ = 100*ms*. (D) Mean squared error between the data and the model for different values of the constant delay time t_0_.

#### 3.2.6 Refining maps by considering constant hemodynamic delays

In section 3.2.3, we have considered that the phase shift solely arise from the low-pass properties of the filter and neglected the constant delay due to sensory integration and signal transmission from the retina to the visual cortex. To incorporate this delay, equation (2) should rather read:

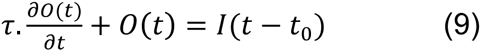

This additional constant phase delay *t*_*0*_ is small, on the order 50-100 ms (Ouellette & Casanova, 2006), but might in fact be crucial for periodic stimuli with short durations or extracted harmonics with high frequencies. We explored this dependence using the data from Fig. 5. For each cycle duration, we plotted the phase shift (Fig. 6C) for the harmonics *H*_*1*_, *H*_*2*_ and *H*_*4*_. The phase Bode plot (τ = 5 *s*, *t*_*0*_ = 0) was superimposed in black, full line. While data were generally distributed around the expected phase shift (Linear regression, R^2^ = 0.677), phase delays extracted at very high pulsations (> 1/10 Hz) typically deviated from the model. We thus took the full model (9) incorporating the constant delay and searched for the optimal value *t*_*0*_ fitting the data (Fig. 6D). We found that the best fit was for *t*_*0*_ = 100 ms, and confirmed that this value greatly improved the constancy with the data (Fig. 6A, black dashed line), particularly for the highest frequencies (R^2^ for *H*_*4*_ data in red increased from 0.365 to 0.731).

### 3.3 APPLICATION TO FAST AND ACCURATE RECONSTRUCTION OF DIRECTION MAPS

We use the methodology developed in the previous sections to reduce the distortions of the different harmonics in order to reconstruct accurate direction maps in cat visual cortex. Following our analysis, we pre-processed the data by inverting the first-order low-pass filter (τ = 5 *s*) before extracting phase and magnitude maps.

The phase maps at *H*_*2*_ reportedly represent the orientation map (Kalatsky and Stryker, 2003; Zepeda et al., 2004; Jha et al., 2005; Yakoub et al., 2008; Vanni et al., 2010a; Sun & al., 2013)).

Similarly, the phase map extracted at *H*_*1*_ (Fig. 7B) was originally proposed to extract the preferred direction map (Kalatsky et al., 2003). Further investigations confirmed this idea, but reported only a mild orthogonal relationship between phase maps extracted at *H*_*1*_ and *H*_*2*_ (Vanni et al., 2010a) contrasting classical episodic methods (Kisvarday et al., 2001; Swindale et al., 2003; Ribot et al., 2008). This is also visible in our data, as shown in Fig. 7D (R^2^ = 0.659). In addition, a prominent topological feature of the direction maps is that they present linear discontinuities, called fractures, across which the preferred direction is reversed. These lines are connected to pinwheel centers in the orientation maps (Tanaka, 1997; Kisvarday et al., 2001; Swindale et al., 2003; Imamura et al., 2006). However, when considering the fractures of the *H*_*1*_ phase map, we found that they were noisy and hardly approached pinwheel centers (Fig. 7C). These properties thus suggest that the phase map extracted at *H*_*1*_ is not sufficient to fully describe the preferred direction map.

**Figure 7.**
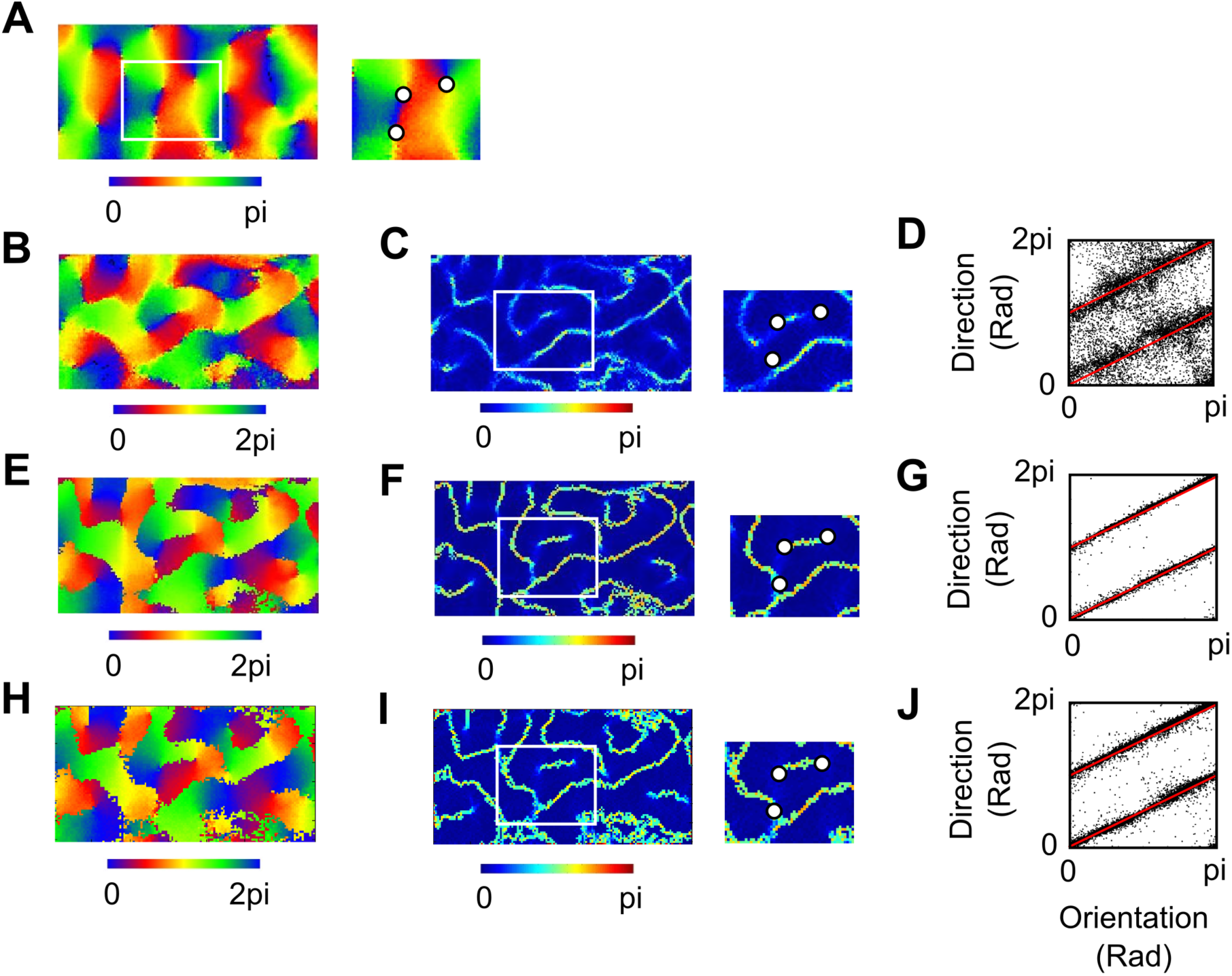
Reconstruction of preferred direction maps. Phase maps were rectified by inverting the first-order low-pass filter with τ = 5 *s* (best actual value τ = 4.66 *s*) (A) Preferred orientation map extracted at *H*_*2*_. Right: A zoom on a domain containing three pinwheels (white dots). (B-D) Properties of the phase map extracted at *H*_*1*_. (B) Phase map extracted at *H*_*1*_. (C) Norm of the direction map gradient. (D) Relationship between the orientation map and the phase map extracted at *H*_*1*_. (E-G) Same as (B-D) when summing up signals extracted at *H*_*1*_ and *H*_*2*_. (H-J) Same as (B-D) for the episodic technique after Von Mises fitting.

To overcome this difficulty, we reconstructed at each pixel the modulation induced by the periodic stimulation from phases and magnitudes extracted at *H*_*1*_ and *H*_*2*_ simultaneously. The preferred direction was then defined as the stimulus direction for which the resulting tuning curve was maximal. The resulting direction map is shown in Fig. 7E. As shown in Fig. 7F-G, this simple method greatly increased the well-known properties of the direction map. This allowed not only to recover efficiently the fractures, but also to extract precisely how these are connected to orientation pinwheel centers (Fig. 7F). Moreover, a strong orthogonality with the orientation map was observed (Fig. 7G., R^2^ = 0.991). These results were comparable with those obtained with episodic methods after Von Mises fitting (Fig. 7H-J, Swindale et al., 2003, R^2^ = 0.961 between the preferred orientation and the preferred direction).

## 4 DISCUSSION

In this study, we observed that hemodynamic signals of functional maps follow a first-order differential equation, resulting in a low-pass temporal filtering of neuronal activity. This filtering has direct consequences on phase delays and signal attenuation in continuous, periodic stimulation paradigms. These findings thus allow recovering accurate functional maps simply by inverting this filter. We illustrated this technique on cat primary visual cortex and showed how to obtain fine orientation and direction maps that do not suffer artifacts of current signal analysis methods. By focusing on the functional hemodynamic response, that are drastically different from HRF arising in response to a single condition, this methodology provides new insight on the dynamics of the hemodynamic signal and new avenues for fast and accurate analysis of functional imaging data.

### 4.1 The hemodynamic signal as a first-order low-pass filter

We have evidenced the filtering properties of the hemodynamic signal on functional maps by presenting stimuli with various durations. Our results argue that the orientation magnitude follows a first-order differential equation representing a low-pass temporal filter. This model can be improved in several directions. We have seen that considering higher-order dynamics do not substantially improve the model and introduces biologically implausible time constants. However, incorporating a constant delay modeling sensory integration and signal transmission from the retina to the visual cortex improved the consistency with the data.

The low-pass nature of the hemodynamic signal was confirmed by highlighting the various phase delays and magnitude decreases when the periodic, continuous stimulation paradigm is used. The results indicate, in particular, that the slope in the magnitude bode diagram for frequencies larger than the cutoff frequency was around −20 dB/decade, which is typical of a first-order low-pass filter.

How this first-order low-pass filter relates to the well-studied HRF (generally modeled as a difference between two alpha functions) in response to an impulse stimulus requires further investigations. A possibility is that the collective transformation of multiple HRFs into functional maps cancels some non-specific activity contained in the HRF. For instance, it is known that even non-optimal stimuli induced a hemodynamic response in the primary visual cortex (Grinvald et al., 1999). This activity is weaker that the activity induced by an optimal stimulus and should be partially suppressed when all the stimulus responses are combined together.

### 4.2 Implications for neuroimaging experiments based on periodic, continuous stimulation

Over the last decade, periodic stimulation paradigms have allowed to efficiently extract functional features in the sensory cortex, to drastically increase the signal to noise ratio, and to shorten the recording time. It has been successfully applied to various cortical areas, including the auditory cortex in rat (Kalatsky et al., 2005; Polley et al. 2007) and in ferret (Nelken et al., 2008), or the somatosensory barrel cortex in mice (Arnett et al., 2014). But this technique has been mostly applied in the primary visual cortex (V1), which is commonly used as a landmark for many studies. It has allowed visualizing various organizations, such as orientation map (Cat: Jha et al., 2005; Kalatsky and Stryker, 2003; Vanni et al., 2009. Tree shrews: Zepeda et al., 2004. Humans: Sun & al., 2013, Yakoub et al., 2008), the retinotopic map (Mouse: Kalatsky and Stryker, 2003; Cat: Vanni et al. 2010b; Macaque: Lu et al., 2009; Human: Engel et al., 1997), the direction map (Vanni et al., 2010a) or the spatial frequency map (Cat: Ribot et al., 2013, 2016; Romagnoni et al., 2015).

We show that the filtering properties of the hemodynamic signal induced stereotyped distortions in the recorded signal when neurons were continuously activated through periodic stimulation. Phase delays were shifted depending on the extracted harmonics, implying that a frequency-dependent rectification should be performed to recover the true phase maps as obtained with conventional techniques. These results strongly differ from the general belief that a common hemodynamic delay is responsible for the shift.

Magnitude maps were also attenuated, in accordance with the impact of a low-pass filter on the signal. Here again, it is important to rectify these activities since they are used for calculating features selectivity: direction (and orientation) selectivity can be estimated by taking the ratio between the magnitude maps of the first and second harmonics (second and fourth harmonics, respectively). Since attenuation is frequency-dependent, a true estimation of the direction (and orientation) selectivity cannot be achieved without a rectification.

Eventually, our data supports that the reconstruction of the preferred direction map with continuous, periodic techniques is possible. In accordance with previous results (Vanni et al., 2010a), the phase map extracted at *H*_*1*_ was not sufficient to fully reproduce the known topological properties exhibited by direction map. We propose instead that phase and magnitude maps extracted at *H*_*1*_ and *H*_*2*_ should be combined. This implies again that both phase delays and magnitude changes induced by the low-pass dynamics of the functional maps should be rectified.

### 4.3 Choice of the parameters for the stimulation and the analysis

The value of the time constant τ (or the cutoff frequency ω_*c*_) is critical to invert the first-order low-pass filter. This value should ideally be defined for each experiment individually since the neurovascular coupling may vary depending on the experimental conditions (Pisauro et al., 2007). In theory, a single value can be readily calculated for each experiment from the phase difference between two harmonics. In practice, however, we found this method to be rather unstable: the stimulation frequency being right into the linear zone of the phase Bode diagram (Fig. 2A), fitting the phase difference between harmonics led to large variations in the estimation of τ (data not shown). We suggest instead to use τ = 5 *s* as a common value among experiments. Indeed, we found a high constancy in the estimation of τ when comparing the phase maps obtained with episodic and continuous methods (5.03 ± 0.55 s.d).

Our results also provide additional information on the impact of the stimulation frequency ω on the recorded data. Pioneering investigations have emphasized that this frequency should be chosen outside the frequency ranges of physiological noise (vasomotor signals, breathing, heart beat…) or of slow variations. Here we show that this choice is also determined by the magnitude attenuation imposed by the hemodynamic filter. In particular, for extracted frequencies ω ≫ ω_*c*_, the magnitude of the signal is weak and the model requires the addition of a constant delay to fully account for the phase shifts. Frequencies ω ≪ ω_*c*_ do not suffer from these issues, but make the recorded time unnecessary long. Appropriate extracted frequencies should thus be adjusted around ω_*c*_ (i.e. ∼ 1/30 Hz with τ = 5 *s*), for which the decrease of magnitude is not too strong, and that do not require the inclusion of a constant time delay in the model to recover the functional maps.

## 5. Conclusion

We developed a novel approach for analyzing data obtained with continuous, periodic stimulation techniques. By revealing the low-pass temporal properties of the hemodynamic signal, we argued that a simple inversion of first-order low-pass filter allowed recovering accurate functional maps, with minimal distortions compared to episodic techniques but massive reduction of the experiment duration. This simple approach strongly differs from common analysis techniques since it does not require inferring the hemodynamic response function, but only the filter time constant, which we showed to be extremely robust across different animals. These findings allowed fast and accurate reconstruction of functional maps from intrinsic signals imaging. We illustrated this technique for direction maps in cat V1. This methodology shall also provide new perspectives for Human functional imaging that rely on hemodynamic signals such as fMRI or diffuse optical imaging.

## Author contributions

A.S., G.H. and J.R. analyzed the data; J.R. designed and performed experiments; A.S., J.T. and J.R. built the model. J.T. and J.R. supervised the study and wrote the paper.

## LEGENDS

**Supplementary Info. 1.**
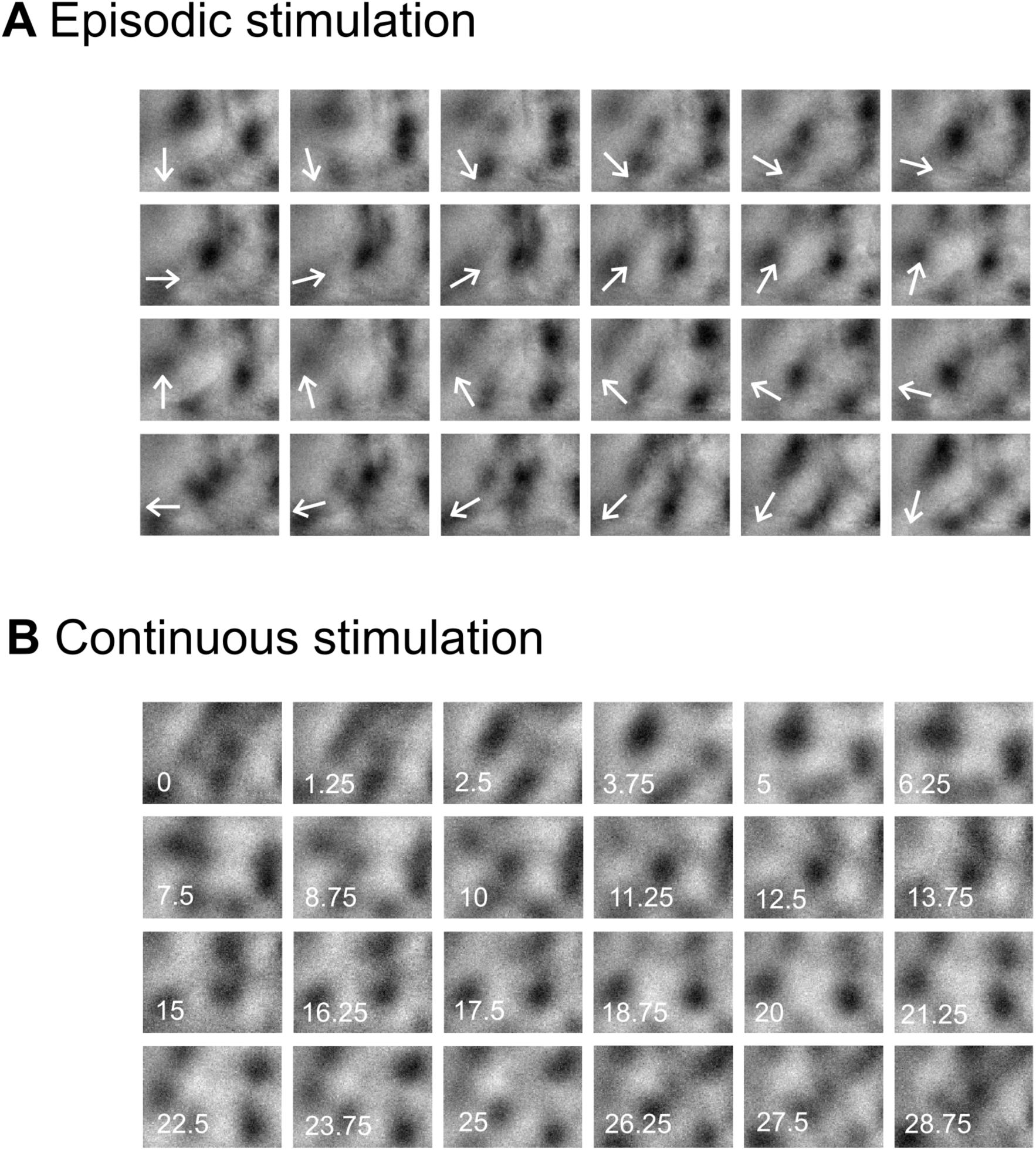
Single condition maps for phase maps in Fig. 4 based on episodic and continuous methods. (A) Single condition maps obtained with the episodic stimulation protocol. Stimuli consisted in oriented gratings that drifted perpendicular to the orientation (N = 24 directions). The direction of motion is indicated with an arrow. Domains in black indicate an activity. (B) Single condition maps obtained with the continuous stimulation protocol. For comparison with (A), each map around 1.25 s are shown, although data were in fact collected each 300 ms (see Methods). Note that these maps resembled best those in (A) after a delay of 4-6 s.

